# Extracellular vesicles promote autophagy in human microglia through lipid raft-dependent mechanisms

**DOI:** 10.1101/2023.07.03.547488

**Authors:** Diana Romenskaja, Ugnė Jonavičė, Augustas Pivoriūnas

## Abstract

Autophagy dysfunction has been closely related with pathogenesis of many neurodegenerative diseases and therefore represents a potential therapeutic target. Extracellular vesicles (EVs) may act as a potent anti-inflammatory agents and also modulators of autophagy in target cells. However, the molecular mechanisms by which EVs modulate autophagy flux in human microglia remain largely unexplored.

In the present study we investigated the effects of EVs derived from human oral mucosa stem cells on the autophagy in human microglia. We demonstrate that EVs promoted autophagy and autophagic flux in human microglia and that this process was dependent on the integrity of lipid rafts. LPS also activated autophagy, but combined treatment with EVs and LPS suppressed autophagy response indicating interference between these signalling pathways. Blockage of Toll-like receptor 4 (TLR4) with anti-TLR4 antibody suppressed EV-induced autophagy. Furthermore, blockage of EV- asscoiated HSP70 chaperone which is one of the endogenous ligands of the TLR4 also suppressed EV- induced lipid raft formation and autophagy. Pre-treatment of microglia with selective inhibitor of αvβ3/αvβ5 integrins cilengitide inhibited EV-induced autophagy. Finally, blockage of purinergic P2X4 receptor (P2X4R) with selective inhibitor 5-BDBD also suppressed of EV-induced autophagy.

In conclusion, we demonstrate that EVs activate autophagy in human microglia through interaction with HSP70/TLR4, αVβ3/αVβ5, and P2X4R signalling pathways and that these effects depend on the integrity of lipid rafts.

Our findings could be used for development of new therapeutic strategies targeting disease-associated microglia.

## Introduction

Microglial cells regulate immune homeostasis in the central nervous system (CNS) by constant monitoring of the brain tissue for signs of damage and clearing cellular debris [1,2]. Abnormal activation of microglia has been associated with many acute and chronic inflammatory CNS disorders, therefore targeting of disease-associated microglia represents a promising therapeutic approach [1,3].

Extracellular vesicles (EVs) are nanosized lipid bilayer – enclosed vesicles, that carry different proteins, RNAs, lipids, and other bioactive molecules [4]. Increasing evidence demonstrate that EVs may act as a potential immunomodulatory and anti-inflammatory agents [5,6]. Moreover, some unique characteristics of EVs such as biocompatibility, specific targeting and accumulation in pathologically affected areas, low immunogenicity and the ability to cross biological barriers make them an attractive therapeutic tool for a variety of neurological disorders [7–10]. Intravenously administered EVs derived from mesenchymal stem cells (MSC) reduced neuroinflammation and switched microglia towards restorative functions after cortical injury in the aged brain of monkeys [11]. EVs administered either by intracerebroventricular, or by intranasal routes, protected mice from the neonatal stroke by direct interaction with microglial cells [12]. EVs also suppressed neuroinflammation and rescued cognitive impairments after traumatic brain injury [13]. However, the mechanisms by which EVs regulate neuroinflammatory response of microglia remain largely unexplored.

Autophagy dysfunction has been closely related with pathogenesis of many neurodegenerative diseases [14]. In this regard, it is important to note that autophagy can also serve as controlling mechanism of the metabolic and immune status of microglia and thus balance neuroinflammatory response [14–16]. For instance, ATG7 gene-deficient microglial cells, show transcriptional and functional similarity to the functionally incompetent microglia from the aged wild-type mice [17]. Induction of autophagy in aged mice using disaccharide trehalose resulted in the restoration of microglial phagocytic activity and remission of the neuroinflammation [17]. Autophagy also negatively regulates inflammasomes in microglia during Aβ-induced neuroinflammation [18] and experimental autoimmune encephalomyelitis [19,20]. Several experimental evidence indicate bidirectional relationship between autophagy and EVs. Autophagy is crucial for the synthesis, secretion and degradation of the EVs. Increased levels of autophagy have been associated with inhibition of EV release due to the increased fusion of multivesicular bodies with autophagic vacuoles [21–23]. On the other hand, exogenous EVs can modulate autophagic flux in target cells. Microglia-derived EVs promoted autophagy and mediated multic:target target signalling during microgliac:microglia crosstalk *in vitro* [24]. Neural stem cell-derived EVs attenuated apoptosis and neuroinflammation after traumatic spinal cord injury by activating autophagy [25]. By contrast, in ischemic stroke model, EVs from human iPSC-derived MSC suppressed autophagy and promoted angiogenesis *in vivo* and *in vitro* [26]. These findings suggest that EVs can be used for targeting autophagy in the disease-associated microglia. However molecular mechanisms by which exogenous EVs modulate autophagic flux in target cells remain unknown.

Our previous findings demonstrate that EVs may act as a potent immunomodulators of human microglia by suppressing TLR4/NFκB signalling pathway, promoting phagocytosis and inducing metabolic reprogramming [5]. More recently we demonstrated that EVs increased migration of human microglia through milk fat globule-epidermal growth factor-VIII (MFG-E8)/ purinergic P2X4R receptor (P2X4R) pathways [27]. EVs also induced lipid raft formation in human microglia through Toll-like receptor 4 (TLR4), P2X4R and integrin αVβ3/αVβ5 signalling pathways [28]. In the present study we investigated the effects of EVs on the autophagy in human microglia. We demonstrate that EVs activate autophagy and autophagic flux in human microglia through TLR4, αVβ3/αVβ5, and P2X4R signalling pathways and that these effects depend on the integrity of lipid rafts.

## Materials and methods

### Human oral mucosal stem cell line

Human oral mucosal stem cells (OMSCs) were isolated from three retromolar explants of a healthy man, under the permission of the Bioethics committee. The biopsy was performed during dental surgery, and the explants were immediately placed in 1 g/l glucose DMEM medium containing 10% FBS and a triple dose of antibiotics. Each explant was transferred to a separate well of a 6-well plate, moistened with 100 μl of low glucose DMEM (Biochrom) (1 g/l), 10% FBS, and 200 U/ml penicillin, 200 μg/ml streptomycin (all from Biochrom, Berlin, Germany), and stored at 37 °C in a humidified incubator with 5% CO_2_. After adhesion of the explants (approximately 3 to 4 hours), each well was filled with medium. Explants were maintained at 37 °C in a humidified atmosphere with 5% CO_2_, and the medium was routinely changed twice per week. After the appearance of migrating cells, the explants were removed from the wells and the medium changed every three days until the cell cultures reached subconfluence.

### Human microglial cell line

The immortalized (SV40) human microglial cell line was acquired from ABM. Human microglial cells were cultured in cell culture flasks coated with 50 μg/ml rat tail collagen I (Gibco) in high glucose DMEM (4.5 g/l) supplemented with GlutaMAX (Gibco) and 10% FBS (Biochrom) depleted of extracellular vesicles (EVs).

### Isolation of extracellular vesicles

EVs were isolated by differential centrifugation according to Thery et al [29]. with some modifications. All centrifugation steps were performed at 4 °C. Supernatants from OMSCs cultured in FBS medium depleted of EVs were successively centrifuged at increasing speeds (300 g for 10 minutes, 2000 g for 10 minutes, then 20000 g for 30 minutes). The final supernatants were ultracentrifuged at 100 000 g for 70 minutes in a Sorvall LYNX 6000 ultracentrifuge with a T29-8×50 rotor in oak ridge centrifuge tubes with caps (all from Thermo Fisher Scientific, Rochester, NY), then the pellets were resuspended in ice-cold PBS and ultracentrifuged again at 100 000 g for 70 minutes at 4°C. The final pellet of EVs (exosomal fraction) was resuspended in filtered ice-cold PBS and stored at -70 °C. Nanoparticle tracking analysis (NTA) was performed using NanoSight LM10 (Malvern Panalytical). The NTA analyses revealed that the EV fractions contained vesicles with a size of approximately 152 nm and Western blot analysis showed that they expressed relatively high amounts of CD63, HSP70 and MFG-E8 proteins (Supplemental figure 1). The yield of EVs obtained from the supernatants of OMSCs grown to subconfluence on cell culture flasks (with an area of 37.5 cm^2^) and conditioned for 72 hours in high-glucose DMEM containing ultracentrifuged EV-depleted FBS medium was defined as one activity unit (1 AU). According to NTA measurements, 1AU of EV contained 3.65 × 10^8^ vesicles.

### Protein isolation and western blot analysis

For cell lysate preparation, cells were washed three times with cold PBS, and lysed in Pierce RIPA buffer supplemented with 1× Halt protease inhibitor cocktail for 15 minutes on ice. Cells were scraped off, and lysates were transferred to eppendorfs for 10 minutes. Samples were centrifuged at 18000 g for 20 minutes at 4°C. After centrifugation, the supernatants were transferred to the new 1.5 ml tubes. For protein isolation from the EVs, we used precipitation with cold acetone. Briefly, four volumes of cold (−20°C) acetone were added to the EV suspension, mixed, and incubated overnight at -20°C. The next day, after centrifugation (20000 g for 15 minutes at 4°C), pellets were washed three times with 80 % acetone using the same conditions. Protein concentrations were measured using the NanoPhotometer Pearl (Implen, Munchen, Germany). For Western blot analysis, cell lysates were diluted in Laemmli sample buffer and heated at 95 °C for 5 minutes. Denatured proteins were separated by sodium dodecyl sulfate-polyacrylamide gel electrophoresis (SDS-PAGE) and blotted onto a PVDF membrane in a semidry Trans-Blot Turbo transfer system (Bio-Rad). Membranes were blocked for 1 hour at room temperature with 5% BSA in PBS containing 0.18% Tween-20 (PBS-Tw). Membranes were then probed overnight at 4°C with primary antibodies against HSP70 (BD Biosciences), CD63, and MFG-E8 (both from Santa Cruz Biotechnology). After washing three times in PBS-Tw, the membranes were incubated with horseradish peroxidase (HRP)-conjugated secondary antibody (Thermo Fisher Scientific) for 1 hour at room temperature. The washing procedure was repeated, and immunoreactive bands were detected with Clarity ECL Western blotting substrate (Bio-Rad) using the ChemiDoc MP system (Bio-Rad).

### Transmission electron microscopy

Transmission electron microscopy (TEM) of EVs was conducted following a previously established protocol [29] with some adjustments. In summary, extracellular vesicles (EVs) suspended in PBS were treated with 2% paraformaldehyde (PFA) for 40 minutes on ice to ensure fixation. Copper grids coated with Formvar and carbon were placed on a 10-µl droplet of the fixed EV suspension at room temperature for 20 minutes. Subsequently, the grids were rinsed with PBS and transferred to a 30-µl droplet of 1% glutaraldehyde solution for 5 minutes at room temperature. The grids underwent eight consecutive washes by transferring them from one droplet of distilled water to another. To enhance contrast, the samples were treated with 2% neutral uranyl acetate on 30 µL droplets for 5 minutes at room temperature in the absence of light. Afterward, the grids were air-dried for 5 minutes. Analysis of the samples was carried out using a FEI Morgagni 268 transmission electron microscope.

### Immunocytochemistry

The autophagy process in microglial cells was evaluated immunocytochemically by staining against autophagy marker LC3B protein [30]. Immediately after treatments, cells were fixed in freshly prepared 4% PFA for 20 minutes at room temperature. Then the cells were washed 3 times with PBS and blocked with 1% BSA-PBS solution for 30 min at room temperature. Afterwards, the cells were incubated at 4 °C overnight with the primary antibodies against LC3B (Sigma-Aldrich) diluted with a blocking solution in a ratio of 1:100. Next day, the cells were washed 3 times with PBS and incubated with the secondary antibodies conjugated with Alexa Fluor 594 (1:1000, Thermo Fisher Scientific) diluted in PBS at room temperature, in the dark, for 1 hour. Then the cells were washed three times with PBS and mounted on slides with Duolink® PLA mounting medium containing DAPI. Prepared samples were analyzed by Leica TCS SP8 confocal microscope (Leica Microsystems, Mannheim, Germany). Images were acquired with a 63× oil immersion objective. Quantification was performed using Leica Application Suite X (LAS X) software (Leica Microsystems). Mean fluorescence intensity values from each field of view were divided by the number of cells detected (determined by counting DAPI-stained nuclei). Data was collected from at least three biological replicates and quantification was performed in at least 15 fields of view for each experimental group.

Alternatively, autophagy and autophagic flux were assessed using RFP-GFP-LC3B chimeric protein. We used Premo™ Autophagy Tandem Sensor RFP-GFP-LC3 Kit (P36239, Molecular Probes, Thermo Fisher Scientific, Bremen, Germany) that allows visualization of the maturation of the autophagosomes to the autolysosomes. This kit contains the BacMam reagent, which is a baculovirus with a mammalian promoter that encodes the tagRFP-eGFP-LC3B chimeric protein. Briefly, the working principle of this reagent is as follows: sequence inserted into the baculovirus vector combine an acid-sensitive GFP with an acid-insensitive RFP, consequently, the change from autophagosome (neutral pH) to autolysosome (with an acidic pH) can be visualized by imaging the specific loss of the GFP fluorescence, leaving only red fluorescence [31].

Autophagic flux analysis in microglial cells was performed according to the manufacturer’s protocol with minimal modifications. 2 × 10^4^ microglia cells were seeded onto coverslips in 4-well plates, cultured overnight and then incubated with 8 μl of BacMam reagent for 48 hours. This volume was calculated using the following formula:

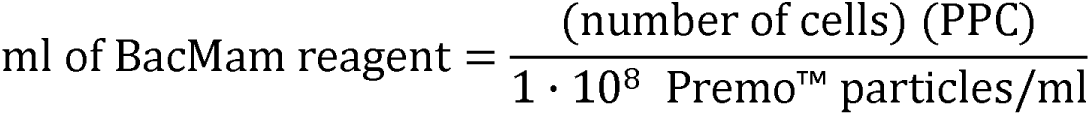

where PPC is the number of viral particles per cell. We tested different PPCs in our microglial cells and found that the highest infection efficiency was achieved when PPC = 40. After 48 hours of incubation with BacMam reagent, cells were exposed to different agents depending on the experimental design. Then samples were prepared and analyzed as described previously, the only difference is that in this case both GFP and RFP fluorescence were analyzed and quantified separately. After specific loss of GFP fluorescence following fusion with lysosomes (Fig.1), calculation of the RFP:GFP ratio was used as an indicator of autophagic flux.

**Figure 1.**
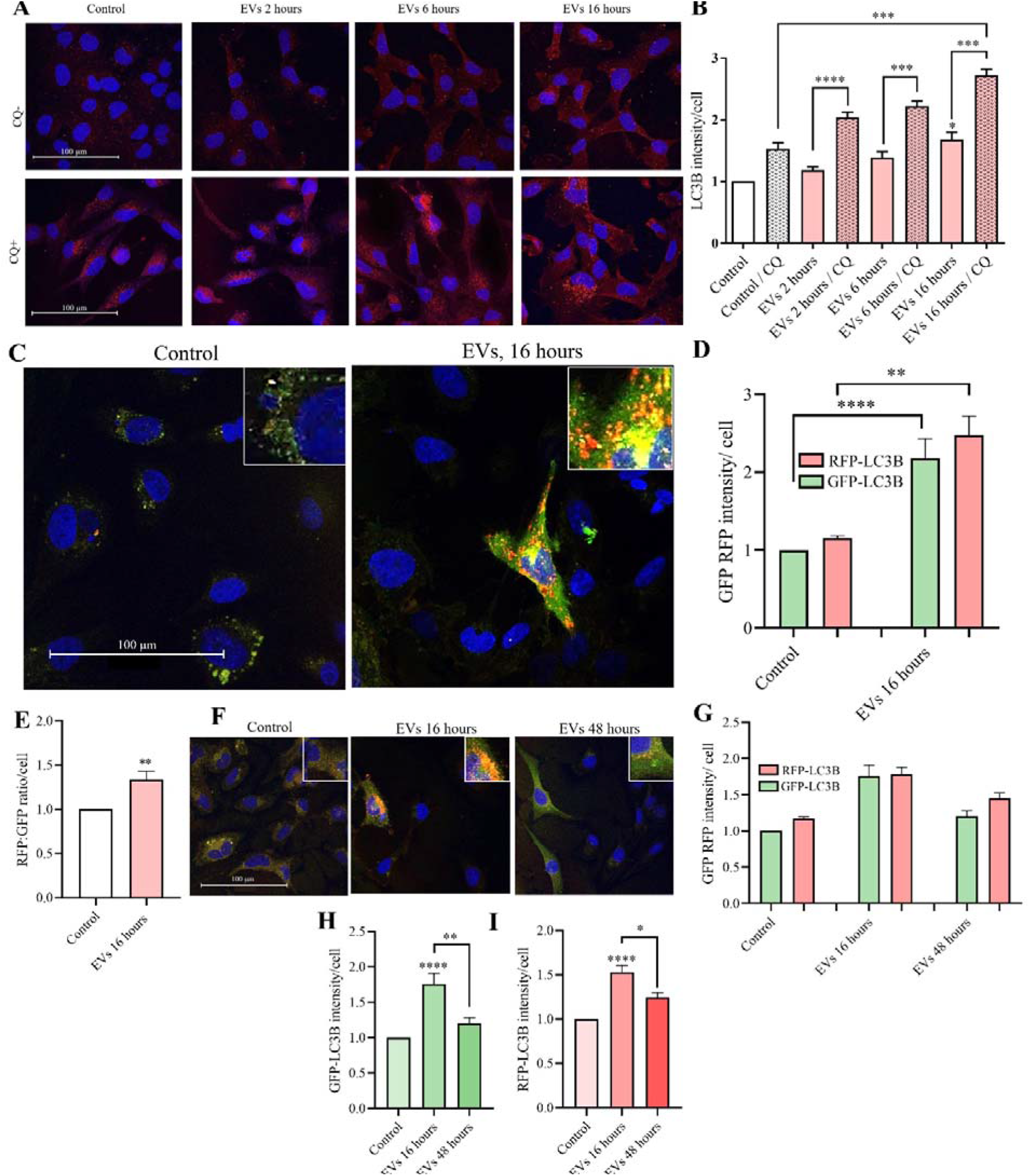
Effects of EVs on autophagy in human microglia. **A** – Confocal images of LC3B protein (red) in microglia cells treated with EVs for different durations (scale bar = 100 μm, magnification – 63x). **B** – Mean fluorescence intensity of LC3B protein per cell measured using LAS X software. Data represent the mean ± SEM from 20 fields of view (n = 4), results normalized to control. Statistical significance was determined using Kruskal-Wallis test followed by Dunn’s post-hoc test in GraphPad Prism 8.0.1 software (* p<0.05; *** p<0.001; **** p<0.0001). **C** – Representative confocal images of microglial cells infected with Premo™ Autophagy Tandem Sensor RFP-GFP-LC3 Kit. **D** – Statistical analysis of GFP-LC3B and RFP-LC3B protein expression. Data shown represent the mean ± SEM from 15 fields of view (n = 3). Statistical significance was determined using Kruskal-Wallis test followed by Dunn’s post-hoc test (** p<0.01). **E** – Estimation of autophagic flux by RFP:GFP ratio. Data represent the mean ± SEM from three independent experiments (n = 3). Statistical significance was determined using the Mann-Whitney U test (** p<0.01). **F** – Confocal images of microglial cells infected with Premo™ Autophagy Tandem Sensor RFP-GFP-LC3 Kit after exposure to EVs for different durations. **G** - Quantification data of GFP-LC3B and RFP-LC3B protein expression. Data shown represent the mean ± SEM from 15 fields of view (n = 3). Statistical analysis was performed using Kruskal-Wallis test followed by Dunn’s post-hoc test for GFP-LC3B (**H**) and RFP-LC3B (**I**) protein expression (* p<0.05; *** p<0.001)

### Experimental design

To assess the effect of EVs on autophagy and autophagic flux, microglial cells were incubated with 1 AU of EVs for 2, 6 and 16 hours, afterwards some samples were additionally treated with 30 μM of chloroquine (CQ) (Sigma-Aldrich, lysosomotropic agent) for 4 hours.

For lipid raft disruption microglial cells were pre-incubated with 5 mM of methyl-β-cyclodextrin (MβCD) (Sigma-Aldrich) for 1 hour.

To assess the effect of LPS (Sigma-Aldrich) on autophagy and autophagic flux microglial cells were incubated with 5 μg/ml of LPS for 16 hours, we also tested the combined treatment of 1 AU of EVs and 5 μg/ml of LPS for 16 hours. For TLR4 blocking cells were pre-incubated with 0.5 μg/ml anti-TLR4 blocking antibody (Abcam, HT125) for 4 hours and then treated with LPS or (and) EVs. Cells were also incubated with 0.5 μg/ml mouse IgG isotype control (Santa Cruz Biotechnology) for 4 hours as a specificity control for anti-TLR4 antibody.

Blockage of EV HSP70 protein was performed by the pre-incubation of EVs with 1 μg/ml anti-HSP70 antibody (Invitrogen) [32] in high glucose DMEM (4.5 g/l) supplemented with GlutaMAX (Gibco) and 10% FBS (Biochrom) depleted of EVs, on the rocker platform for 2 hours at 37°C. As a specificity control for anti-HSP70 antibody we used 1 μg/ml rabbit IgG isotype control (Invitrogen) for 2 hours.

For inhibition of P2X4R cells were pre-incubated for 30 min with 10 μM of 5-BDBD (Tocris Bioscience). Blockage of αvβ3/αvβ5 integrins was achieved by the pre-incubation with 10 μM of cilengitide for 2 hours (Sigma-Aldrich).

### Lipid raft labeling

Lipid raft labeling was performed with Vybrant® Alexa Fluor® 594 Lipid Raft Labeling Kit (Thermo Fisher Scientific) according to the manufacturer’s protocol with minimal modifications. After all treatments, cells were washed with complete growth medium and incubated with 1 μg/ml of fluorescent CT-B (cholera toxin subunit B) for 10 minutes at room temperature. CT-B selectively binds to ganglioside GM1 specifically enriched in the lipid rafts. After washing three times with PBS, the fluorescent CT-B-labeled lipid rafts were cross-linked with the anti-CT-B antibody (200-fold dilution in complete growth medium) for 15 minutes at room temperature. Cells were then washed three times with PBS and fixed in freshly prepared 4% PFA for 20 minutes at room temperature. Cells were washed three times with PBS and mounted on slides with Duolink® PLA mounting medium containing DAPI and analyzed using a Leica TCS SP8 confocal microscope (Leica Microsystems, Mannheim, Germany). Images were acquired with a 63× oil immersion objective. Quantification was performed using Leica Application Suite X (LAS X) software (Leica Microsystems). Mean fluorescence intensity values from each field of view were divided by the number of cells detected (determined by counting DAPI-stained nuclei). Data was collected from three biological replicates and quantification was performed in 15 fields of view for each experimental group.

### Statistical analysis

Statistical analysis was performed using data from at least three independent biological experiments. Plots show the mean and standard error of the mean (SEM). Since all our data did not meet the assumptions of parametric methods (failed the Shapiro-Wilk normality test or variances were unequal), nonparametric tests were used for statistical analysis. Difference between two groups was determined using the Mann-Whitney U test, and differences between 3 or more groups were determined by Kruskal-Wallis one-way analysis of variance followed by Dunn’s post-hoc test. All results were considered significant when p < 0.05. Data was analyzed using Graph Pad Prism® 8.0.1 version software (Graph Pad Software, Inc., San Diego, CA, USA).

## Results

### EVs promote autophagy and autophagic flux in human microglia

We first tested how OMSC-derived EVs affect LC3B expression in human microglial cells. For this purpose, the cells were incubated with 1 AU of EVs for 2, 6 and 16 hours (Fig.1 A CQ-). The change in autophagic flux was determined by comparing the amount of autophagosomes in the absence of a lysosomal inhibitor and after inhibition of autophagosome degradation with chloroquine (CQ) (Fig.1 A CQ+). CQ inhibits the fusion of autophagosomes with lysosomes, accumulating in lysosomes as a deprotonated weak base, thereby increasing lysosomal pH [33]. Our results show that EVs significantly increased LC3B protein expression after 16 hours of treatment (compared to control by 67.7 %, n = 4, p = 0.0131; Fig. 1 B). We observed the same effects when the cells were additionally treated with CQ. The highest expression of LC3B was detected after 16 hours (compared to control/CQ by 78.5 %, n = 4, p = 0.0002, Fig. 1 B). EVs also promoted autophagic flux in human microglial cells. Exposure to EVs for 2, 6 and 16 hours increased autophagic flux by 73.1%, 59. 8% and 62.3%, respectively (n = 4, p _2h_ < 0.0001, p _6h_ = 0.0007, p _16h_ = 0.0002; Fig. 1 B). Based on these results, an optimal exposure of 16 hours was chosen for further experiments.

These results were further confirmed with RFP-GFP-LC3B reporters (Premo™ Autophagy Tandem Sensor Kit). Exposure to EVs for 16 hours significantly increased autophagosome formation (p = 0.0135, n = 3, Fig. 1 D) and autophagic flux (p = 0.0049, n = 3, Fig. 1 E) in microglia cells. We also found that 48 hours after exposure to EVs autophagy response declined to the basal levels (p_GFP-LC3B_ = 0.0142, p_RFP-LC3B_ = 0.0297, Fig.1 F, G, H, I).

### Disruption of lipid rafts prevents EV-induced autophagy in microglia

We have previously demonstrated that EVs promoted lipid raft formation in human microglia [27,28]. We therefore asked whether EV-induced autophagy depends on the integrity of lipid rafts. For this purpose cells were pre-treated for 1 hour with 5 mM cholesterol removing agent methyl-β-cyclodextrin (MβCD) and then exposed to the 1 AU of EVs for 16 hours (Fig. 2).

**Figure 2.**
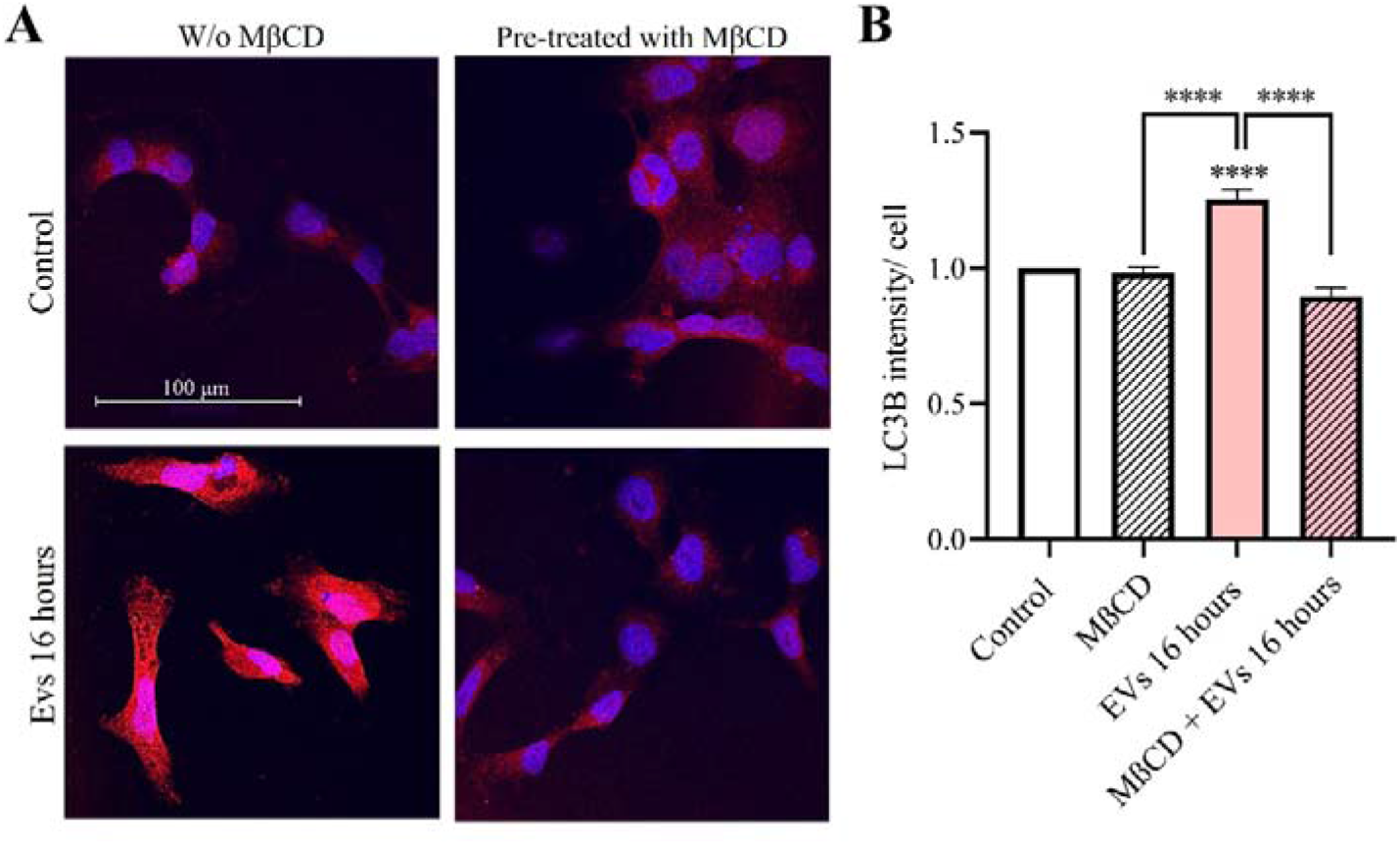
Autophagy activation assay after removing one of the main components of lipid rafts. **A** – Confocal images of labeled LC3B protein (red) in microglia cells in the presence and absence of pre-treatment with 5 mM MβCD for 1 hour (63× magnification immersion objective, scale bar = 100 μm). **B** – The mean fluorescence intensity of LC3B protein per cell were measured with Leica Application Suite X (LAS X) software, data shown represent the results of 15 fields of view for each experimental group from three independent biological experiments (n = 3), plotted as the mean ± SEM, results normalized to control. Statistical significance was analyzed by Kruskal–Wallis test followed by Dunn’s post-hoc test in GraphPad Prism 8.0.1 software (**** p < 0.0001)

Disruption of lipid raft suppressed EV-induced autophagy to the basal levels (by 28.6%, p <0.0001, n = 3; Fig. 2 B). Our results demonstrate that lipid raft integrity is essential for the EV-induced autophagy in human microglia.

### Combined treatment with EVs and LPS attenuate autophagy response in human microglia

We have recently demonstrated that combined treatment with EVs and LPS decreased lipid raft formation in microglia [28]. We therefore tested how combined treatment with EVs and LPS affect autophagy response. Microglial cells were exposed to EVs (1 AU), or LPS (5 µg/ml) and combination of both EVs and LPS for 16 hours (Fig. 3).

**Figure 3.**
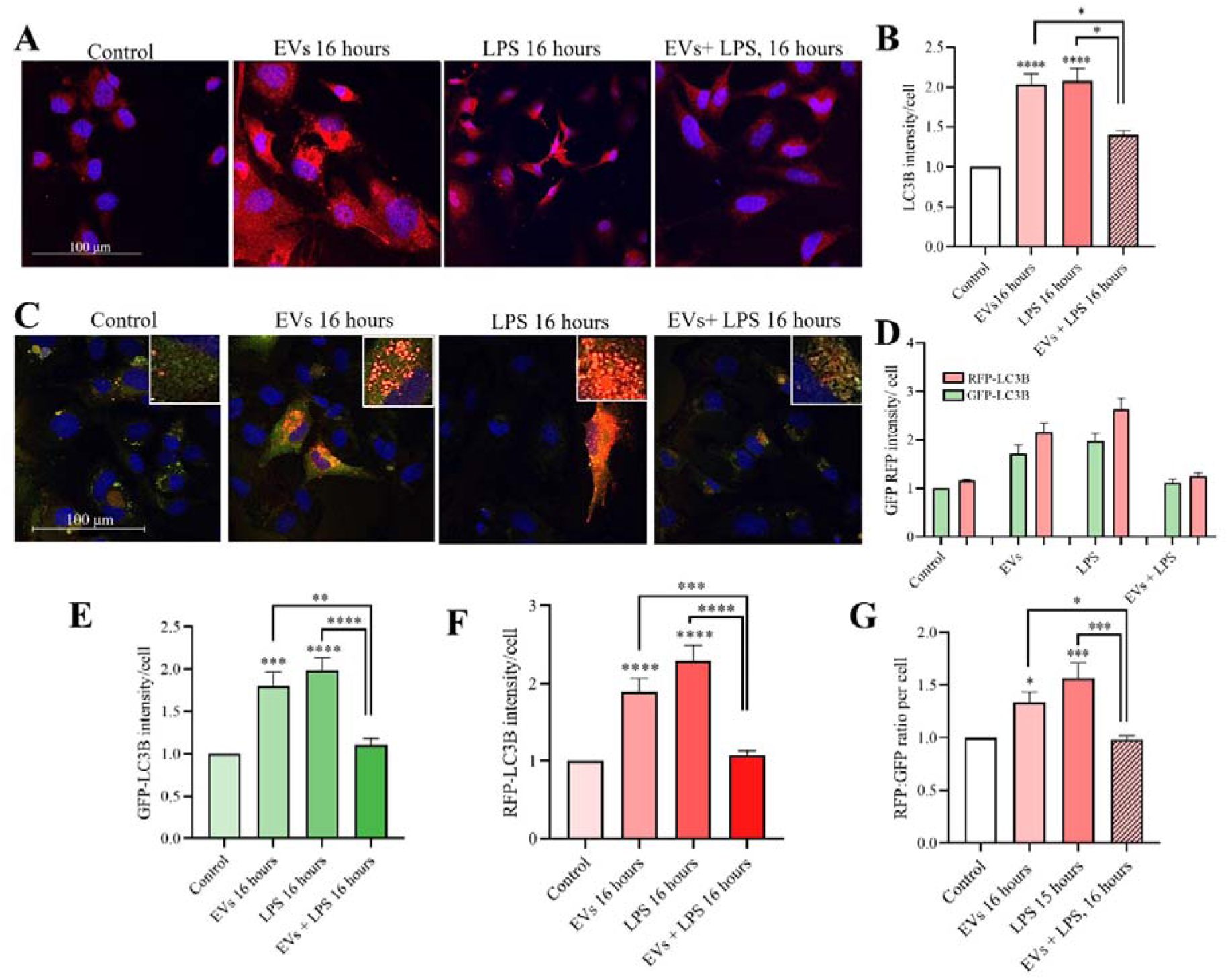
Effects of EVs and LPS on autophagy in human microglia cells. **A** – Confocal images of LC3B protein (red) in microglia cells treated with EVs (1 AU) and LPS (5 μg/ml) for 16 hours (scale bar = 100 μm, magnification – 63x). **B** – Mean fluorescence intensity of LC3B protein per cell measured using LAS X software. Data represents the mean ± SEM from 15 fields of view (n = 3). Statistical significance was determined using Kruskal-Wallis test followed by Dunn’s post-hoc test in GraphPad Prism 8.0.1 software (* p<0.05; **** p<0.0001). **C** – Confocal images of microglial cells infected with Premo™ Autophagy Tandem Sensor RFP-GFP-LC3 Kit. **D** – Quantification data of GFP-LC3B and RFP-LC3B protein expression. Data shown represents the mean ± SEM from 15 fields of view (n = 3). Statistical analysis was performed using Kruskal-Wallis test followed by Dunn’s post-hoc test for GFP-LC3B (**E**) and RFP-LC3B (**F**) protein expression (* p<0.05; ** p<0.01; *** p<0.001). **G** – Estimation of autophagic flux by RFP:GFP ratio. Data represents the mean ± SEM from three independent experiments (n = 3). Statistical significance was determined using Kruskal-Wallis test followed by Dunn’s post-hoc test (* p<0.05; *** p<0.001).

We found that both EVs and LPS similarly increased autophagy (EVs by 103.6%, p <0.0001; LPS by 107.6%, p <0.0001, n = 3, Fig. 3 A, B). By contrast, combined treatment with EVs and LPS suppressed autophagy response (by 32.7% when compared to LPS, p = 0.0327, and by 31.3% when compared to EVs, p = 0.0411, n = 3; Fig. 3 A, B).

We next tested how combined treatment with EVs and LPS affect autophagy and autophagy flux in microglia expressing tagRFP-eGFP-LC3B reporters (Fig. 4 C). We found that both EVs and LPS significantly increased signal intensity of GFP-LC3B protein (EVs by 94.5 %, p = 0.0202 and LPS by 93.1%, p = 0.0059, n = 3, Fig. 3 E). By contrast, combined treatment suppressed GFP-LC3B signal to the control levels (Fig.3 E). These results indicate that when used separately, both EVs and LPS promote autophagosome formation, whereas combined treatment suppress autophagosome formation. Similar results were obtained with RFP-LC3B protein. Exposure to EVs increased signal intensity of RFP-LC3B protein by 88.3 % (p = 0.0105, n = 3, Fig. 3 F) and to LPS by 157 % (p = 0.0002, n = 3, Fig. 3 F), whereas combined treatment suppressed signal to the control levels. Calculations of the RFP:GFP ratios which represent the activity of the autophagic flux showed that when used separately both EVs and LPS activate autophagic flux (EVs – by 33.7 %, p = 0.0421 n = 3, LPS – by 56.6 %, p = 0.0004, n = 3, Fig. 3 G). Again, combined treatment with EVs and LPS suppressed autophagy flux to the control levels (Fig. 3 G).

Our findings show mutual interference between EVs and LPS signalling during induction of autophagy in human microglia.

**Figure 4.**
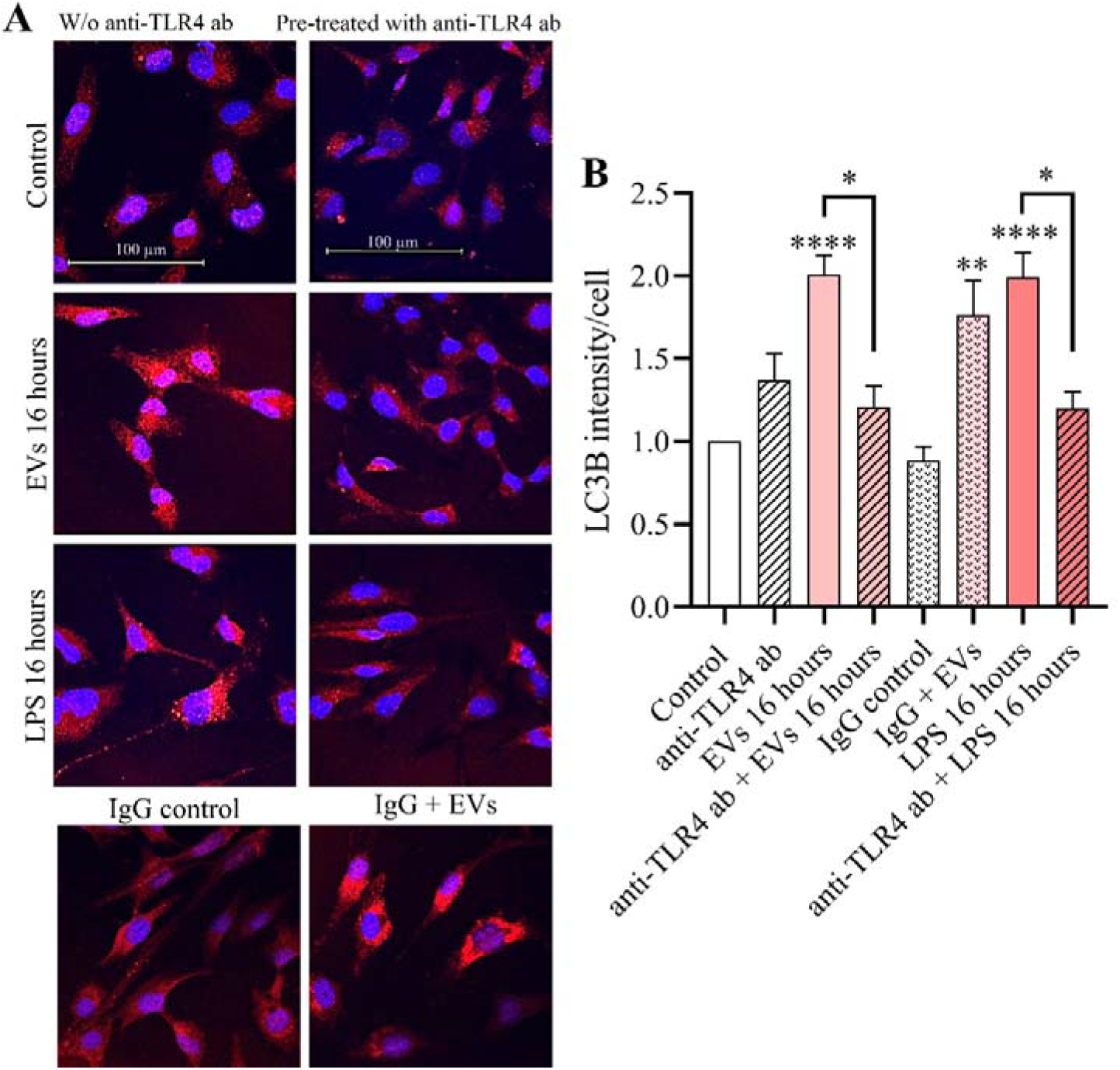
Anti-TLR4 antibody prevents EVs-induced effect. **A** – confocal images of labeled LC3B protein (red) in microglia cells. The cells were treated with EVs (1 AU) and LPS (5 μg/ml) for 16 hours before immunocytochemistry and fixation (scale bar = 100 μm, magnification – 63x). **B** – the mean fluorescence intensity of LC3B protein per cell were measured with Leica Application Suite X (LAS X) software. Data shown represent the results of 15 fields of view for each experimental group from three independent biological experiments (n = 3), plotted as the mean ± SEM, results normalized to control. Statistical significance was analyzed by Kruskal–Wallis test followed by Dunn’s post-hoc test in GraphPad Prism 8.0.1 software (* p<0.05; ** p<0.01; **** p < 0.0001)

### Blockage of TLR4 prevents EV-induced autophagy

We further investigated possible crosstalk between EVs and LPS signalling in microglia and tested the effects of specific blockage of TLR4 on the EV-induced autophagy (Fig. 4).

As expected, pre-treatment with anti-TLR4 antibody prevented LPS-induced autophagy (p = 0.0295, n = 3, Fig. 4 A, B). Importantly, blockage of TLR4 also suppressed EVs-induced autophagy in microglia (decreased by 39.9 %, p = 0.0119, n = 3; Fig. 4 B). In contrast, pre-treatment with control IgG antibody did not affect EV-induced autophagy. Our results demonstrate that EVs trigger autophagy response in microglia through interaction with TLR4.

### EV-associated HSP70 chaperones promote formation of lipid rafts and autophagy in human microglia

HSP70 protein is one of the endogenous ligands of the TLR4 [34]. HSP70/TLR4 interaction is also important for the protective effect of exosomes against neomycin-induced hair cell death [32]. We detected high expression levels of HSP70 in our EV preparations (Supplemental figure 1 C). We therefore used neutralizing anti-HSP70 antibody for blockage of the EV-associated HSP70 and then investigated lipid raft formation and autophagy in human microglia cells.

Pre-treatment of EVs with a neutralizing anti-HSP70 antibody (1 μg/ml) for 2 hours resulted in a significant decrease in their ability to induce lipid raft formation in microglial cells (p = 0.0009, n = 3, Fig. 5 B). Pre-treatment with IgG control antibody did not affect EV-induced lipid raft formation. We also found that blockage of vesicular HSP70 significantly suppressed EV-induced autophagy (p_GFP-LC3B_ = <0.0001, p_RFP-LC3B_ = 0.0005, n = 3, Fig. 5 C, D, E, F) whereas control IgG antibody did not.

**Figure 5.**
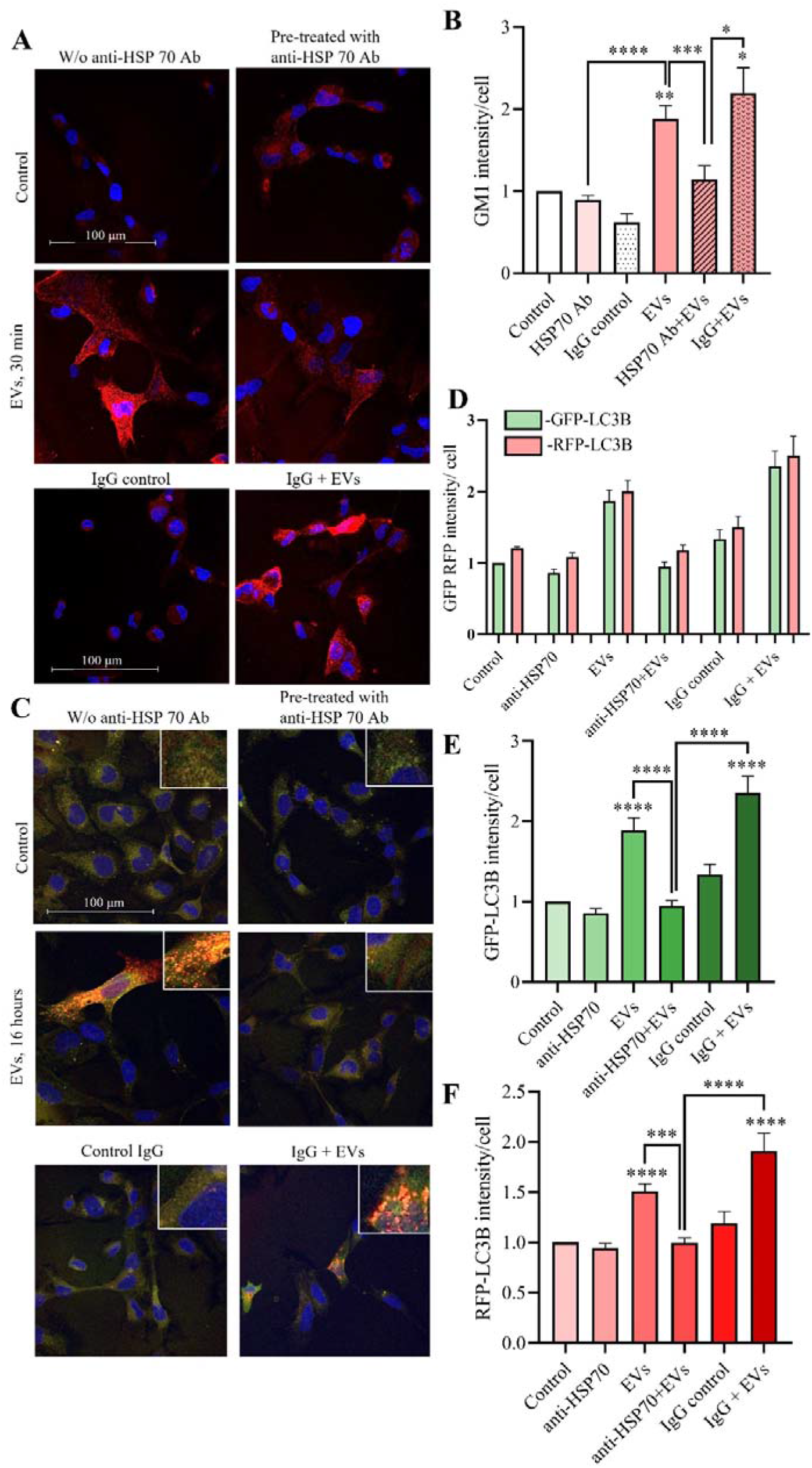
Blockage of EV-associated HSP70 suppresses lipid raft formation and autophagy. **A** – Confocal images of labeled lipid rafts (red) in human microglia cells treated with a selective P2X4R antagonist 5-BDBD (scale bar = 100 μm, magnification – 63x). **B** – Mean fluorescence intensity of labeled ganglioside GM1 per cell measured using LAS X software. Data represents the mean ± SEM from 15 fields of view (n = 3), results normalized to control. Statistical significance was determined using Kruskal-Wallis test followed by Dunn’s post-hoc test in GraphPad Prism 8.0.1 software (*** p<0.001; **** p<0.0001). **C** – Confocal images of microglial cells infected with Premo™ Autophagy Tandem Sensor RFP-GFP-LC3 Kit. **D** – Quantification data of GFP-LC3B and RFP-LC3B protein expression. Data shown represents the mean ± SEM from 15 fields of view (n = 3). Statistical analysis was performed using Kruskal-Wallis test followed by Dunn’s post-hoc test for GFP-LC3B (**E**) and RFP-LC3B (**F**) protein expression (* p<0.05; ** p<0.01; *** p<0.001).

Our results show, that EV-associated HSP70 may act as an endogenous ligand of TLR4 promoting lipid raft formation and autophagy in human microglia.

### Blockage of **α**v**β**3 and **α**v**β**5 integrins prevents EV-induced autophagy in microglia

We have previously demonstrated that EVs express high levels of secretory glycoprotein MFG-E8 (milk fat globule EGF factor) (Supplemental figure 1) [27,28]. MFG-E8 protein can act as molecular bridge by interacting with the cellular αvβ3 and αvβ5 integrins and phosphatidylserine (PS) exposed on the EV membranes. (Fig.7 A) [35,36]. Our recent data demonstrate that blockage of αvβ3 and αvβ5 integrins with cilengitide prevented EV-induced lipid raft formation in microglia cells [28]. We therefore tested the effects of cilengitide on the EV-induced autophagy in microglia (Fig. 6 B, C).

**Figure 6.**
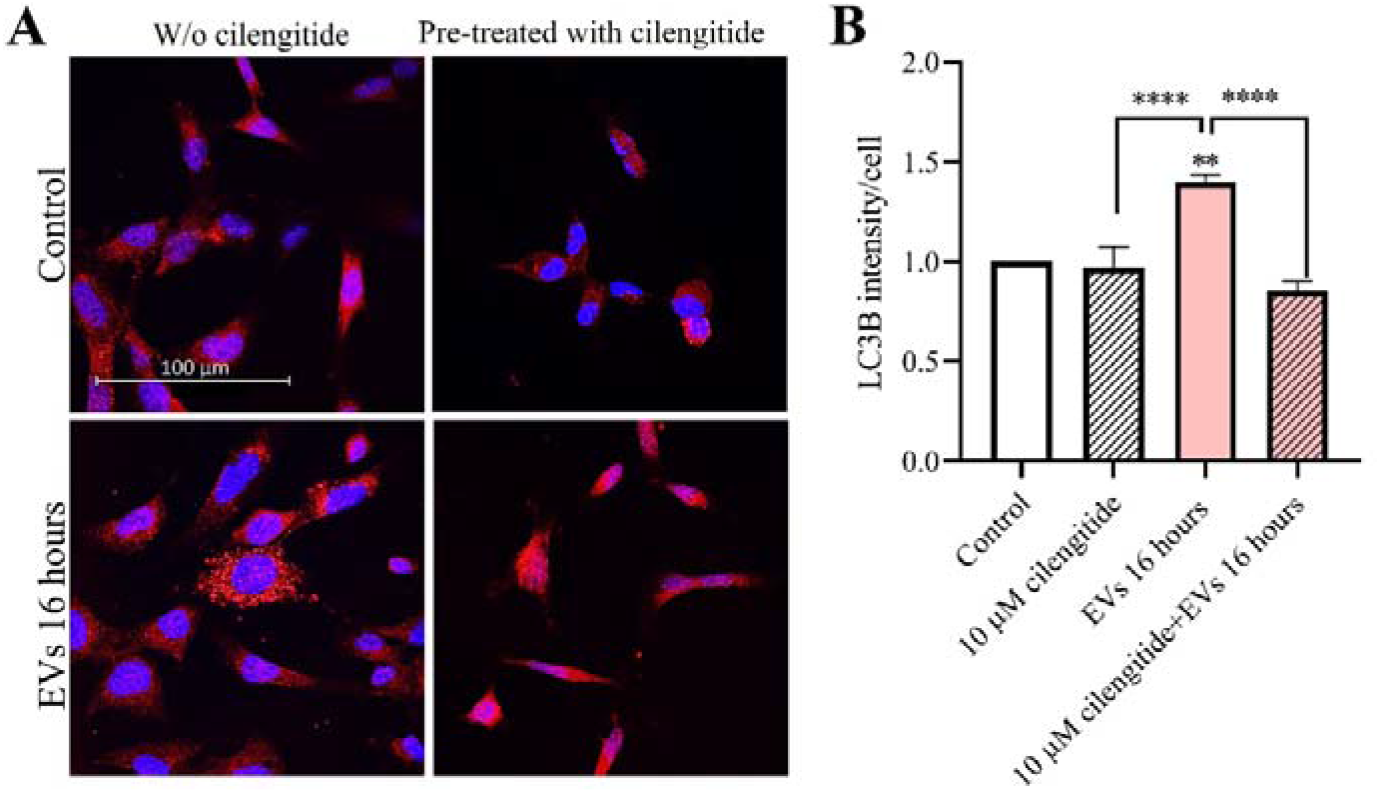
The role of αvβ3/αvβ5 integrins in EVs-induced autophagy. **A** – Confocal images of labeled LC3B protein (red) in microglia cells in the presence and absence of pre-treatment with 10 μM of cilengitide for 2 hours (63× magnification immersion objective, scale bar = 100 μm). **B** – The mean fluorescence intensity of LC3B protein per cell were measured with Leica Application Suite X (LAS X) software, data shown represent the results of 15 fields of view for each experimental group from three independent biological experiments (n = 3), plotted as the mean ± SEM, results normalized to control. Statistical significance was analyzed by Kruskal–Wallis test followed by Dunn’s post-hoc test in GraphPad Prism 8.0.1 software (** p < 0.01; **** p < 0.0001)

**Figure 7.**
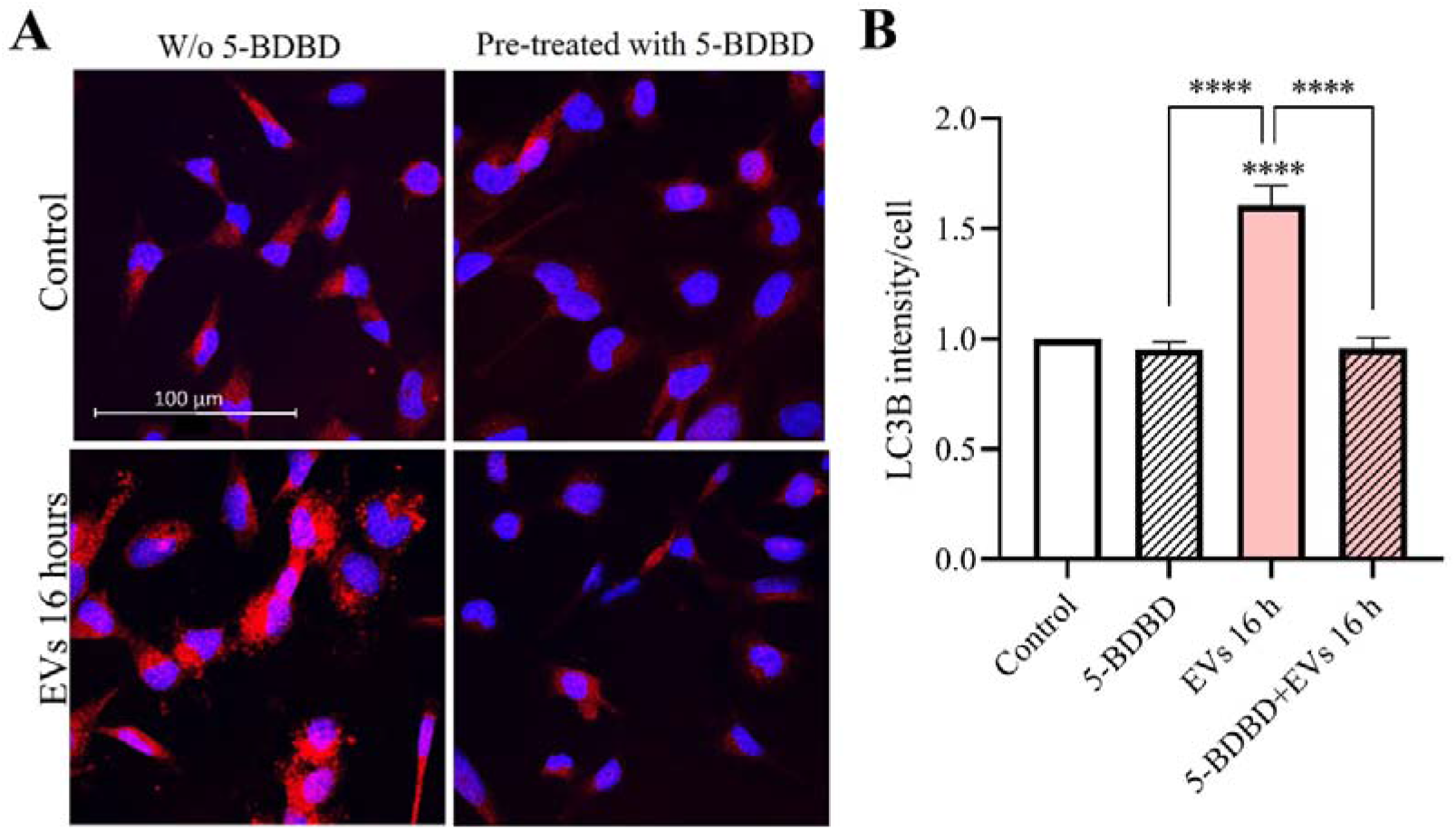
Involvement of P2X4 receptor in EV-induced autophagy. **A** – Confocal images of labeled LC3B protein (red) in microglia cells in the presence and absence of pre-treatment with 10 μM 5-BDBD for 30 min (63× magnification immersion objective, scale bar = 100 μm). **B** – The mean fluorescence intensity of LC3B protein per cell were measured with Leica Application Suite X (LAS X) software, data shown represent the results of 15 fields of view for each experimental group from three independent biological experiments (n = 3), plotted as the mean ± SEM, results normalized to control. Statistical significance was analyzed by Kruskal–Wallis test followed by Dunn’s post-hoc test in GraphPad Prism 8.0.1 software (**** p < 0.0001)

We found that pre-treatment of microglia with 10 μM of cilengitide for 2 hours suppressed EV-induced LC3B protein expression to the basal levels (n = 3, p <0.0001, Fig. 6 C). These findings indicate that αvβ3 and αvβ5 integrins are indispensable for the EV- induced autophagy in microglia.

### Inhibition of P2X4 receptor suppresses EVs-induced autophagy

Our previous study showed a close association between MFG-E8 protein and P2X4 receptor, which increased after exposure to EVs [27]. In this study we tested how blockage of P2X4R with selective antagonist 5-BDBD affect EV-induced increase of LC3B protein in human microglia cells (Fig. 7 A).

Importantly, pre-treatment with 5-BDBD (10 μM) for 30 min completely suppressed EV – induced autophagy (n = 3, p <0.0001, Fig. 7 B) in microglia.

## Discussion

The primary goal of this study was to examine the effects of OMSC-derived EVs on the autophagy process in human microglia cells. We demonstrate that EVs significantly increased autophagy and autophagic flux and therefore could be potentially used for targeting autophagy in the disease-associated microglia.

It has been shown that activation of autophagy can promote neuroprotective properties of microglia during neuroinflammation whereas inhibition of this process can lead to the increased neurodegeneration [37–39]. On the other hand, over-enhancement of autophagy may exert adverse effects on cells, leading to the cell death [40,41]. It is therefore important to understand molecular mechanisms responsible for the autophagy-promoting effects of EVs in human microglia.

We have previously demonstrated that EVs promoted lipid raft formation in human microglia [27,28]. In this study we show that EV-induced autophagy depends on the integrity of the lipid rafts. We also demonstrate that when used separately, both EVs and LPS increased autophagy, but combined treatment suppressed autophagy response. Blockage of LPS receptor TLR4 with anti-TLR4 antibody prevented EV-induced autophagy in human microglia. Furthermore, blockage of HSP70 chaperone which is highly expressed in our EV preparations and is one of the endogenous ligands of the TLR4 also suppressed EV-induced lipid raft formation and autophagy. We therefore suggest that EVs can at least partially trigger lipid raft formation and autophagy response in microglia through interaction with TLR4s. Autophagy promotes elimination of intracellular pathogens and is closely associated with different innate immunity signalling pathways [42,43]. Different types of TLR ligands enhanced autophagy in immune cells [44]. In particular, stimulation of TLR4s activated autophagy by affecting Bcl-2 - Beclin 1 interactions in macrophages [45]. Interestingly, several studies emphasize a close interplay between lipid rafts and autophagy machinery. For instance, it has been shown that lipid rafts can regulate autophagy by interacting with autophagosomes and autophagy-related proteins, such as ATG5 and ATG12 [46]. However, the exact molecular mechanisms linking EV/TLR4 – induced lipid raft formation with activation of autophagy in human microglia remain unclear.

Microglia are highly motile cells and we previously showed, that EV-induced migration of microglial cells depends on the αvβ3 and αvβ5 integrins [27]. We also demonstrated that αvβ3/ αvβ5 integrin signalling pathway was indispensable for EV- and LPS-induced lipid raft formation in microglia [28]. In the present study we demonstrate that pre-treatment of microglia with specific inhibitor of αvβ3 and αvβ5 integrins cilengitide suppressed EV-induced autophagy. EV preparations used in our study were highly enriched with MFG-E8 protein (Supplemental figure 1) which can act as a molecular bridge connecting phosphatidylserine (PS) exposed on the outer membranes of the EVs with microglial integrin αVβ3/αVβ5 receptors [47]. We therefore hypothesize that EV-associated MFG-E8 protein may interact with microglial αVβ3/αVβ5 receptors and promote downstream signalling events leading to the increased migration and autophagy. It has been shown that autophagy regulates integrin-mediated cell adhesion and may promote cell migration affecting the turnover of focal adhesions [48,49]. Interestingly, integrins may also play an important role in the autophagosome formation by direct interactions with LC3 proteins [50,51]. We speculate that EVs can affect bidirectional interplay between integrin recycling and autophagy thereby promoting remodeling of focal adhesions and microglial motility.

We have recently demonstrated that pharmacological inhibition of purinergic receptor P2X4R signalling suppressed EV-induced lipid raft formation, migration and phagocytic activity of microglia [27,28]. P2X4Rs undergo rapid and constitutive endocytosis from the plasma membrane to the lysosomes and are activated by both intraluminal ATP and by a decrease in acidity [52]. Activated P2X4Rs initiate release of Ca^2+^ from the lysosomes and promote fusion of the lysosome membranes with the autophagosomes stimulating autophagic flux [53]. In the present study we show that blockage of P2X4R with selective antagonist 5-BDBD inhibited EV – induced autophagy in microglia. Our results are in agreement with other studies showing that P2X4R signalling promotes autophagy activation in different experimental models [54,55]. It is presently unclear, however, whether EVs can directly interact with P2X4Rs on the plasma membrane or induce P2X4R-dependent effects indirectly. Purinergic receptors, TLRs and integrins are enriched in the lipid rafts and therefore are in close proximity [56–58], therefore possible EVs may also directly interact with P2X4Rs. Further studies are needed to elucidate interactions between EVs and P2XR4 signalling pathway.

Another interesting observation of this study is functional interdependence between TLR4, αVβ3/αVβ5, and P2X4R signalling pathways as autophagy promoters in response to the treatment with EVs (Fig. 8 A). We have recently demonstrated that these signalling pathways are functionally interdependent promoters of EV- and LPS- induced lipid raft formation in microglia [28]. We therefore propose that TLR4, αVβ3/αVβ5 integrin, and P2X4R comprise a network of functionally interdependent signalling units regulating the induction and enlargement of lipid rafts leading to the autophagy initiation (Fig. 8).

**Figure 8.**
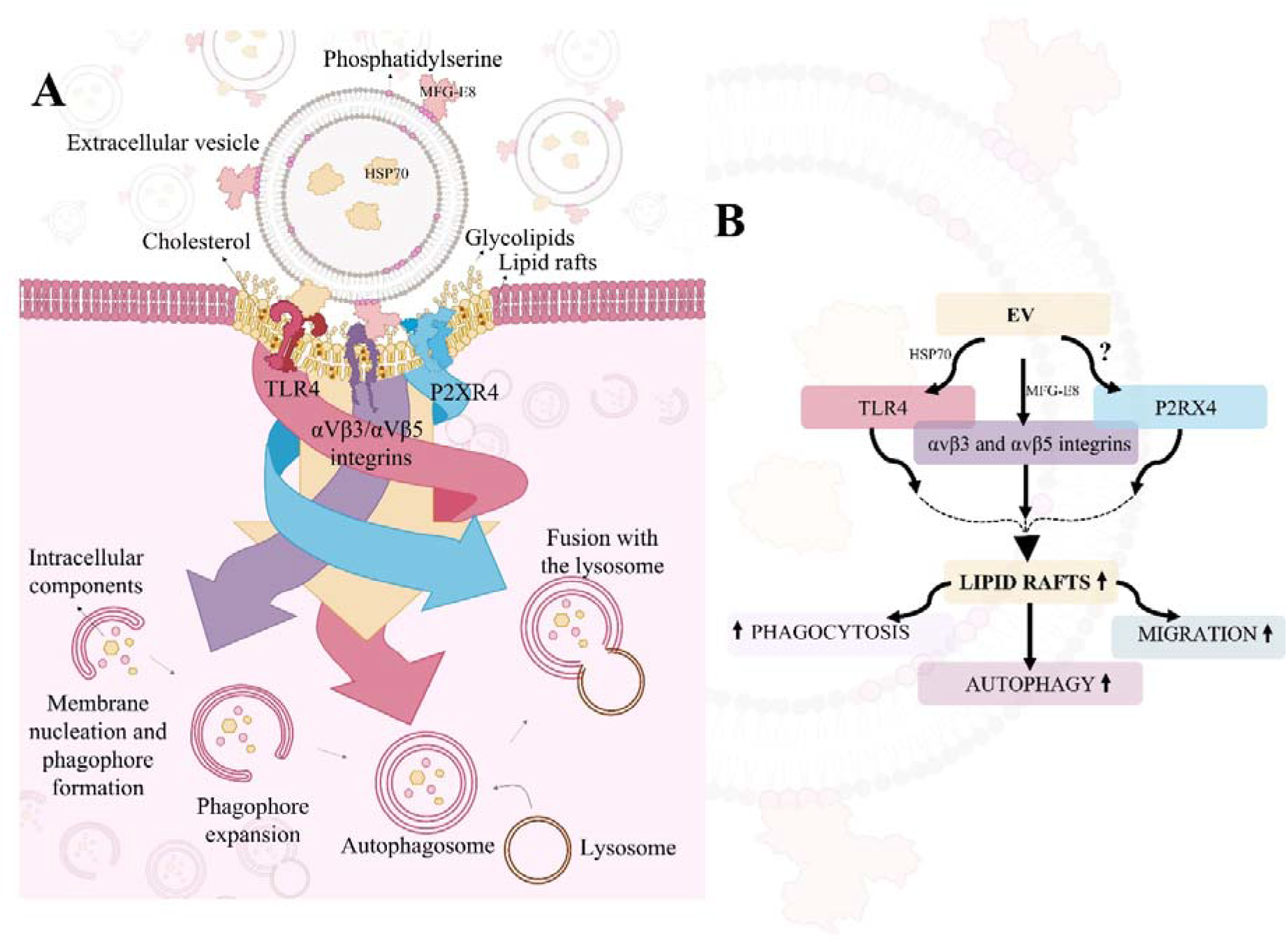
Proposed mechanism for EV action on microglial cells. **A** – An illustration showing a model in which TLR4, αVβ3/αVβ5 integrins, and P2XR4, which localize and activate within lipid rafts, collectively form a network of functionally interdependent signalling units that triggers autophagy in microglia cells. **B** – Results-based scheme showing the effects of EVs on microglial cells

In conclusion, we demonstrate that EVs activate autophagy in human microglia through interaction with TLR4/HSP70, αVβ3/αVβ5, and P2X4R signalling pathways and that these effects depend on the integrity of lipid rafts. Our findings could be used for development of new therapeutic strategies involving extracellular vesicles for targeting disease-associated microglia.

## Acknowledgements

This project has received funding from European Regional Development Fund (project No 01.2.2-LMT-K-718-01-0012) under grant agreement with the Research Council of Lithuania (LMTLT).

## References

1 Augusto-Oliveira M, Arrifano GP, Delage CI, Tremblay M-È, Crespo-Lopez ME & Verkhratsky A (2022) Plasticity of microglia. Biol Rev 97, 217–250.

2 Paolicelli RC, Sierra A, Stevens B, Tremblay ME, Aguzzi A, Ajami B, Amit I, Audinat E, Bechmann I, Bennett M, Bennett F, Bessis A, Biber K, Bilbo S, Blurton-Jones M, Boddeke E, Brites D, Brône B, Brown GC, Butovsky O, Carson MJ, Castellano B, Colonna M, Cowley SA, Cunningham C, Davalos D, De Jager PL, de Strooper B, Denes A, Eggen BJL, Eyo U, Galea E, Garel S, Ginhoux F, Glass CK, Gokce O, Gomez-Nicola D, González B, Gordon S, Graeber MB, Greenhalgh AD, Gressens P, Greter M, Gutmann DH, Haass C, Heneka MT, Heppner FL, Hong S, Hume DA, Jung S, Kettenmann H, Kipnis J, Koyama R, Lemke G, Lynch M, Majewska A, Malcangio M, Malm T, Mancuso R, Masuda T, Matteoli M, McColl BW, Miron VE, Molofsky AV, Monje M, Mracsko E, Nadjar A, Neher JJ, Neniskyte U, Neumann H, Noda M, Peng B, Peri F, Perry VH, Popovich PG, Pridans C, Priller J, Prinz M, Ragozzino D, Ransohoff RM, Salter MW, Schaefer A, Schafer DP, Schwartz M, Simons M, Smith CJ, Streit WJ, Tay TL, Tsai LH, Verkhratsky A, von Bernhardi R, Wake H, Wittamer V, Wolf SA, Wu LJ & Wyss-Coray T (2022) Microglia states and nomenclature: A field at its crossroads. Neuron 110, 3458–3483.

3 Muzio L, Viotti A & Martino G (2021) Microglia in Neuroinflammation and Neurodegeneration. Front Neurosci 15.

4 Wiklander OPB, Brennan M, Lötvall J, Breakefield XO & Andaloussi SEL (2019) Advances in therapeutic applications of extracellular vesicles. Sci Transl Med 11.

5 Jonavičė U, Tunaitis V, Kriaučiūnaitė K, Jarmalavičiūtė A & Pivoriunas A (2019) Extracellular vesicles can act as a potent immunomodulators of human microglial cells. J Tissue Eng Regen Med 13.

6 Čebatariūnienė A, Kriaučiūnaitė K, Prunskaitė J, Tunaitis V & Pivoriūnas A (2019) Extracellular Vesicles Suppress Basal and Lipopolysaccharide-Induced NFκB Activity in Human Periodontal Ligament Stem Cells. Stem Cells Dev 28, 1037–1049.

7 Perets N, Betzer O, Shapira R, Brenstein S, Angel A, Sadan T, Ashery U, Popovtzer R & Offen D (2019) Golden Exosomes Selectively Target Brain Pathologies in Neurodegenerative and Neurodevelopmental Disorders. Nano Lett 19, 3422–3431.

8 Pauwels MJ, Xie J, Ceroi A, Balusu S, Castelein J, Van Wonterghem E, Van Imschoot G, Ward A, Menheniott TR, Gustafsson O, Combes F, EL Andaloussi S, Sanders NN, Mäger I, Van Hoecke L & Vandenbroucke RE (2022) Choroid plexus-derived extracellular vesicles exhibit brain targeting characteristics. Biomaterials 290, 121830.

9 Narbute K, Piļipenko V, Pupure J, Dzirkale Z, Jonavičė U, Tunaitis V, Kriaučiūnaitė K, Jarmalavičiūtė A, Jansone B, Kluša V & Pivoriūnas A (2019) Intranasal Administration of Extracellular Vesicles Derived from Human Teeth Stem Cells Improves Motor Symptoms and Normalizes Tyrosine Hydroxylase Expression in the Substantia Nigra and Striatum of the 6-Hydroxydopamine-Treated Rats. Stem Cells Transl Med 8, 490– 499.

10 Jarmalavičiūtė A, Tunaitis V, Pivoraitė U, Venalis A & Pivoriūnas A (2015) Exosomes from dental pulp stem cells rescue human dopaminergic neurons from 6-hydroxy dopamine-induced apoptosis. Cytotherapy 17, 932–939.

11 Go V, Bowley BGE, Pessina MA, Zhang ZG, Chopp M, Finklestein SP, Rosene DL, Medalla M, Buller B & Moore TL (2020) Extracellular vesicles from mesenchymal stem cells reduce microglial-mediated neuroinflammation after cortical injury in aged Rhesus monkeys. GeroScience 42, 1–17.

12 Pathipati P, Lecuyer M, Faustino J, Strivelli J, Phinney DG & Vexler ZS (2021) Mesenchymal Stem Cell (MSC)–Derived Extracellular Vesicles Protect from Neonatal Stroke by Interacting with Microglial Cells. Neurotherapeutics 18, 1939–1952.

13 Kim D, Nishida H, An SY, Shetty AK, Bartosh TJ & Prockop DJ (2016) Chromatographically isolated CD63+CD81+ extracellular vesicles from mesenchymal stromal cells rescue cognitive impairments after TBI. Proc Natl Acad Sci U S A 113, 170–175.

14 Su P, Zhang J, Wang D, Zhao F, Cao Z, Aschner M & Luo W (2016) The role of autophagy in modulation of neuroinflammation in microglia. Neuroscience 319, 155– 167.

15 Xu Y, Propson NE, Du S, Xiong W & Zheng H (2021) Autophagy deficiency modulates microglial lipid homeostasis and aggravates tau pathology and spreading. Proc Natl Acad Sci 118, e2023418118.

16 Zhu R, Luo Y, Li S & Wang Z (2022) The role of microglial autophagy in Parkinson’s disease. Front Aging Neurosci 14.

17 Berglund R, Guerreiro-Cacais AO, Adzemovic MZ, Zeitelhofer M, Lund H, Ewing E, Ruhrmann S, Nutma E, Parsa R, Thessen-Hedreul M, Amor S, Harris RA, Olsson T & Jagodic M (2020) Microglial autophagy-associated phagocytosis is essential for recovery from neuroinflammation. Sci Immunol 5.

18 Cho M-H, Cho K, Kang H-J, Jeon E-Y, Kim H-S, Kwon H-J, Kim H-M, Kim D-H & Yoon S-Y (2014) Autophagy in microglia degrades extracellular β-amyloid fibrils and regulates the NLRP3 inflammasome. Autophagy 10, 1761–1775.

19 Shao BZ, Wei W, Ke P, Xu ZQ, Zhou JX & Liu C (2014) Activating cannabinoid receptor 2 alleviates pathogenesis of experimental autoimmune encephalomyelitis via activation of autophagy and inhibiting NLRP3 inflammasome. CNS Neurosci Ther 20, 1021–1028.

20 Plaza-Zabala A, Sierra-Torre V & Sierra A (2017) Autophagy and Microglia: Novel Partners in Neurodegeneration and Aging. Int J Mol Sci 18.

21 Wei W, Pan Y, Yang X, Chen Z, Heng Y, Yang B, Pu M, Zuo J, Lai Z, Tang Y & Xin W (2022) The Emerging Role of the Interaction of Extracellular Vesicle and Autophagy— Novel Insights into Neurological Disorders. J Inflamm Res 15, 3395–3407.

22 Xing H, Tan J, Miao Y, Lv Y & Zhang Q (2021) Crosstalk between exosomes and autophagy: A review of molecular mechanisms and therapies. J Cell Mol Med 25, 2297– 2308.

23 Leidal AM & Debnath J (2021) Emerging roles for the autophagy machinery in extracellular vesicle biogenesis and secretion. FASEB BioAdvances 3, 377–386.

24 Van den Broek B, Pintelon I, Hamad I, Kessels S, Haidar M, Hellings N, Hendriks JJA, Kleinewietfeld M, Brône B, Timmerman V, Timmermans JP, Somers V, Michiels L & Irobi J (2020) Microglial derived extracellular vesicles activate autophagy and mediate multi-target signaling to maintain cellular homeostasis. J Extracell Vesicles 10.

25 Rong Y, Liu W, Wang J, Fan J, Luo Y, Li L, Kong F, Chen J, Tang P & Cai W (2019) Neural stem cell-derived small extracellular vesicles attenuate apoptosis and neuroinflammation after traumatic spinal cord injury by activating autophagy. Cell Death Dis 10, 340.

26 Xia Y, Ling X, Hu G, Zhu Q, Zhang J, Li Q, Zhao B, Wang Y & Deng Z (2020) Small extracellular vesicles secreted by human iPSC-derived MSC enhance angiogenesis through inhibiting STAT3-dependent autophagy in ischemic stroke. Stem Cell Res Ther 11, 313.

27 Jonavičė U, Romenskaja D, Kriaučiūnaitė K, Jarmalavičiūtė A, Pajarskienė J, Kašėta V, Tunaitis V, Malm T, Giniatullin R & Pivoriūnas A (2021) Extracellular Vesicles from Human Teeth Stem Cells Trigger ATP Release and Promote Migration of Human Microglia through P2X4 Receptor/MFG-E8-Dependent Mechanisms. Int J Mol Sci 22.

28. 28 Romenskaja D, Jonavičė U, Tunaitis V & Pivoriūnas A (2022) Extracellular vesicles promote lipid raft formation in human microglia through TLR4, P2X4R, and αVβ3/ αVβ5 signaling pathways. *Res Sq*.

29 Théry C, Amigorena S, Raposo G & Clayton A (2006) Isolation and Characterization of Exosomes from Cell Culture Supernatants and Biological Fluids. Curr Protoc Cell Biol 30, 1–29.

30 Baeken MW, Weckmann K, Diefenthäler P, Schulte J, Yusifli K, Moosmann B, Behl C & Hajieva P (2020) Novel Insights into the Cellular Localization and Regulation of the Autophagosomal Proteins LC3A, LC3B and LC3C. Cells 9.

31 Invitrogen Premo ^TM^ Autophagy Tandem Sensor RFP-GFP-LC3B Kit., 1–7.

32 Breglio AM, May LA, Barzik M, Welsh NC, Francis SP, Costain TQ, Wang L, Anderson DE, Petralia RS, Wang YX, Friedman TB, Wood MJA & Cunningham LL (2020) Exosomes mediate sensory hair cell protection in the inner ear. J Clin Invest 130, 2657– 2672.

33 Chen PM, Gombart ZJ & Chen JW (2011) Chloroquine treatment of ARPE-19 cells leads to lysosome dilation and intracellular lipid accumulation: Possible implications of lysosomal dysfunction in macular degeneration. Cell Biosci 1, 1–10.

34 Zhang Y, Zhang X, Shan P, Hunt CR, Pandita TK & Lee PJ (2013) A Protective Hsp70– TLR4 Pathway in Lethal Oxidant Lung Injury. J Immunol 191, 1393–1403.

35 Mai J, Wang K, Liu C, Xiong S & Xie Q (2022) αvβ3-targeted sEVs for efficient intracellular delivery of proteins using MFG-E8. BMC Biotechnol 22, 1–10.

36 Ye H, Li B, Subramanian V, Choi BH, Liang Y, Harikishore A, Chakraborty G, Baek K & Yoon HS (2013) NMR solution structure of C2 domain of MFG-E8 and insights into its molecular recognition with phosphatidylserine. Biochim Biophys Acta - Biomembr 1828, 1083–1093.

37 Jin MM, Wang F, Qi D, Liu WW, Gu C, Mao CJ, Yang YP, Zhao Z, Hu LF & Liu CF (2018) A Critical Role of Autophagy in Regulating Microglia Polarization in Neurodegeneration. Front Aging Neurosci 10, 1–13.

38 Hegdekar N, Sarkar C, Bustos S, Ritzel RM, Hanscom M, Ravishankar P, Philkana D, Wu J, Loane DJ & Lipinski MM (2023) Inhibition of autophagy in microglia and macrophages exacerbates innate immune responses and worsens brain injury outcomes. Autophagy 00, 1–19.

39 Zubova SG, Suvorova II & Karpenko MN (2022) Macrophage and microglia polarization: Focus on autophagy-dependent reprogramming. Front Biosci - Sch 14.

40 Draf C, Wyrick T, Chavez E, Pak K, Kurabi A, Leichtle A, Dazert S & Ryan AF (2021) A Screen of Autophagy Compounds Implicates the Proteasome in Mammalian Aminoglycoside-Induced Hair Cell Damage. Front Cell Dev Biol 9, 1–12.

41 Kar R, Riquelme MA, Hua R & Jiang JX (2019) Glucocorticoid-Induced Autophagy Protects Osteocytes Against Oxidative Stress Through Activation of MAPK/ERK Signaling. JBMR Plus 3, 1–6.

42 Jo EK, Yuk JM, Shin DM & Sasakawa C (2013) Roles of autophagy in elimination of intracellular bacterial pathogens. Front Immunol 4, 1–9.

43 Kuo C-J, Hansen M & Troemel E (2018) Autophagy and innate immunity: Insights from invertebrate model organisms. Autophagy 14, 233–242.

44 Delgado MA, Elmaoued RA, Davis AS, Kyei G & Deretic V (2008) Toll-like receptors control autophagy. EMBO J 27, 1110–1121.

45 Delgado MA & Deretic V (2009) Toll-like receptors in control of immunological autophagy. Cell Death Differ 16, 976–983.

46 Fecchi K, Anticoli S, Peruzzu D, Iessi E, Gagliardi MC, Matarrese P & Ruggieri A (2020) Coronavirus Interplay With Lipid Rafts and Autophagy Unveils Promising Therapeutic Targets. Front Microbiol 11.

47 Cheyuo C, Aziz M & Wang P (2019) Neurogenesis in Neurodegenerative Diseases: Role of MFG-E8. Front Neurosci 13, 569.

48 Vlahakis A & Debnath J (2017) The Interconnections between Autophagy and Integrin Mediated Cell Adhesion. J Mol Biol 429, 515–530.

49 Kenific CM, Wittmann T & Debnath J (2016) Autophagy in adhesion and migration. J Cell Sci 129, 3685–3693.

50 Yang J (2023) Viruses Binding to Host Receptors Interacts with Autophagy. Int J Mol Sci 24.

51 Kliche J, Kuss H, Ali M & Ivarsson Y (2021) Cytoplasmic short linear motifs in ACE2 and integrin β3 link SARS-CoV-2 host cell receptors to mediators of endocytosis and autophagy. Sci Signal 14.

52 Murrell-Lagnado RD & Frick M (2019) P2X4 and lysosome fusion. Curr Opin Pharmacol 47, 126–132.

53 Huang P, Zou Y, Zhong XZ, Cao Q, Zhao K, Zhu MX, Murrell-Lagnado R & Dong X-P (2014) P2X4 forms functional ATP-activated cation channels on lysosomal membranes regulated by luminal pH. J Biol Chem 289, 17658–17667.

54 Chadet S, Allard J, Brisson L, Lopez-Charcas O, Lemoine R, Heraud A, Lerondel S, Guibon R, Fromont G, Le Pape A, Angoulvant D, Jiang L-H, Murrell-Lagnado R & Roger S (2022) P2x4 receptor promotes mammary cancer progression by sustaining autophagy and associated mesenchymal transition. Oncogene 41, 2920–2931.

55 Li R, Lu Y, Zhang Q, Liu W, Yang R, Jiao J, Liu J, Gao G & Yang H (2022) Piperine promotes autophagy flux by P2RX4 activation in SNCA/α-synuclein-induced Parkinson disease model. Autophagy 18, 559–575.

56 Garcia-Marcos M, Dehaye J-P & Marino A (2009) Membrane compartments and purinergic signalling: the role of plasma membrane microdomains in the modulation of P2XR-mediated signalling. FEBS J 276, 330–340.

57 Płóciennikowska A, Hromada-Judycka A, Borzęcka K & Kwiatkowska K (2015) Co-operation of TLR4 and raft proteins in LPS-induced pro-inflammatory signaling. Cell Mol Life Sci 72, 557–581.

58 Leitinger B & Hogg N (2002) The involvement of lipid rafts in the regulation of integrin function. J Cell Sci 115, 963–972.

