## Supplemental figure 1. for "Extracellular vesicles promote autophagy in human microglia through lipid raft-dependent mechanisms"

**Supplementary materials**

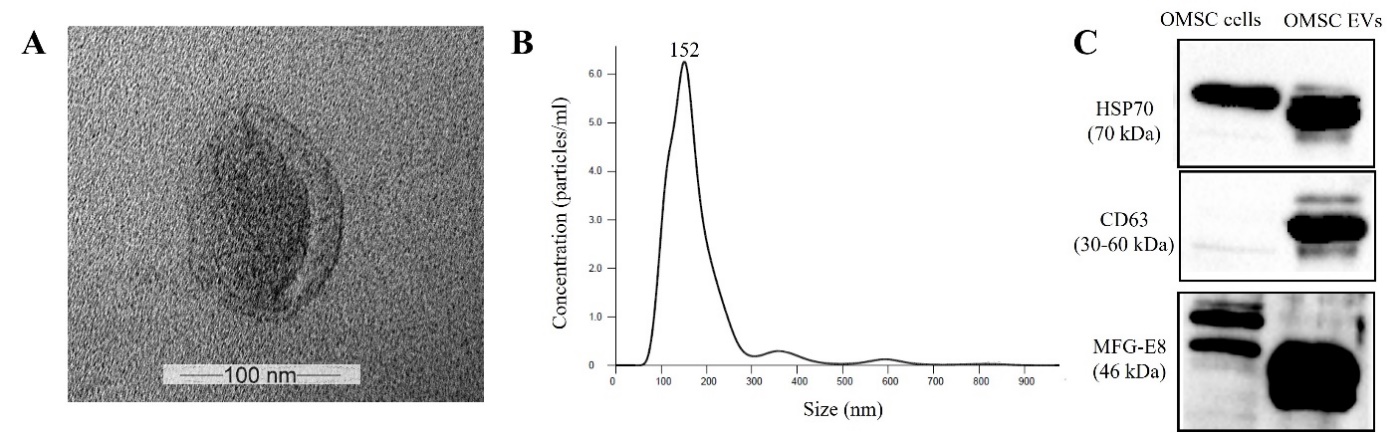

**Supplemental figure 1.** Characterization of extracellular vesicles (EVs) isolated from human oral mucosa stem cells. **A** – Transmission electron microscopy of OMSCs-derived EVs (×100000 magnification). **B** – The concentration and particle size of EVs were analyzed using the NanoSight LM10 instrument (Malvern Panalytical) by nanoparticle tracking analysis. The size distribution of the EVs was approximately 152 nm. **C** – The OMSC cell lysates and OMSC-derived EVs samples underwent electrophoresis, followed by blotting. The resulting membrane was then probed with antibodies targeting HSP70, CD63, and MFG-E8. To visualize the bands, the membrane was incubated with suitable horseradish peroxidase-conjugated secondary antibodies and a chemiluminescence substrate.
